# Integrated microRNA and proteome analysis of cancer datasets with MoPC

**DOI:** 10.1101/2022.09.20.508638

**Authors:** Marta Lovino, Elisa Ficarra, Loredana Martignetti

## Abstract

MicroRNAs (miRNAs) are small molecules that play an essential role in regulating gene expression by post-transcriptional gene silencing. Their study is crucial in revealing the fundamental processes underlying pathologies and, in particular, cancer. To date, most studies on miRNA regulation consider the effect of specific miRNAs on specific target mRNAs, providing wet-lab validation. However, few tools have been developed to explain the miRNA-mediated regulation at the protein level. In this paper, the MoPc computational tool is presented, that relies on the partial correlation between mRNAs and proteins conditioned on the miRNA expression to predict miRNA-target interactions in multi-omic datasets. MoPc returns the list of significant miRNA-target interactions and plot the significant correlations on the heatmap in which the miRNAs and targets are ordered by the chromosomal location. The software was applied on three TCGA/CPTAC datasets (breast, glioblastoma, and lung cancer), returning enriched results in three independent targets databases.

**Author summary:** According to the central dogma of molecular biology, DNA is transcribed into RNA and subsequently translated into proteins. However, many molecules affect the amount of protein produced, including microRNAs (miRNAs). They can inhibit the translation or intervene by implementing the decay of target mRNAs. In literature, most works focus on describing the effect of miRNAs on mRNA targets, while only a few tools integrate protein expression profiles. MoPc predicts miRNA-targets interaction by considering the expression of mRNA, proteins, and miRNAs simultaneously. The method is based on the partial correlation measure between mRNAs and proteins conditioned by the expression of the miRNAs. The results on TCGA/CPTAC datasets prove the relevance of the MoPc method both from a computational and a biological point of view.

## Introduction

MicroRNAs (miRNAs) are small RNA molecules that emerged as important regulators of gene expression at the post-transcriptional level. They are involved in many diverse biological processes [1], and their dysregulation may lead to a variety of diseases, including cancer [2, 3]. Accurate prediction of miRNA-target interactions is critical to understanding the function of miRNAs. Although much progress has been made, target identification remains a challenge because of the limited understanding of the molecular basis of miRNA-target coupling, but also due to the context-dependence of post-transcriptional regulation and miRNA mode of action [4]. Generally, miRNAs downregulate proteins through a combination of translational inhibition and promotion of mRNA decay [5–7], even though which mechanisms of action of microRNAs are the most dominant remains a matter of debate. The emergence of high-throughput methods in the past decades has allowed researchers to address the question of miRNA action on a global scale. High-throughput experiments have usually been focused on measuring miRNA-mediated changes at the mRNA level (degradation), allowing for characterization of only a subset of direct targets [8–11]. Further integration of global proteome profiling has allowed the investigation of miRNA-mediated gene expression changes at both the mRNA and the protein level [12, 13], even though many of these studies are limited to one or a few miRNAs or proteins. Additionally, few published tools are available to integrate mRNA and protein profiles to infer miRNA regulation. ProteoMirExpress [14] infers active miRNAs based on the anticorrelation between miRNAs and their potential targets at either the mRNA or the protein level, or both. However, pairwise correlation approaches applied to omics profiles produce very dense networks containing direct and indirect associations that are very difficult to distinguish.

Here, we propose a novel in silico method, called MoPc (Multi-omics Partial Correlation analysis), to analyze post-transcriptional regulation by miRNAs by integrating information at both mRNA and protein level. The MoPc method predicts miRNA-target potential interactions using high-throughput mRNA, protein, and miRNA expression datasets from matched samples. It computes the conditional dependence (partial correlation) of mRNA and protein levels, taking into account the effect of a third variable, the expression of a given regulatory miRNA. the underlying hypothesis assumes that if a miRNA regulates a given gene, then the partial correlation between the mRNA and the protein expression of that gene conditioned by the expression of the miRNA will be greater than the bivariate correlation between the mRNA and protein levels of the gene. Therefore, to identify candidate miRNA targets, MoCP selects those genes whose partial correlation between the level of RNA and protein levels conditioned to the expression of a given miRNA is more significant than the bivariate correlation between the RNA and the protein.

MoPc has been applied to TCGA/CPTAC omics datasets of breast, glioblastoma, and lung cancer, predicting key miRNAs involved in these diseases. To assess the reliability of the interactions predicted by the MoPc, they have been tested for the enrichment in predicted targets by three independent databases, namely miRDB [15, 16], TargetScan [17], and miRTarBase [18]. The targets predicted by MoPC in breast and lung cancer show significant overlap with those predicted by the three independent databases. In contrast, the targets predicted in glioblastoma significantly overlap with targets predicted in TargetScan and miRTarBase. Finally, MoPc provides the user with a heatmap to visualize results and simultaneously explore the conditional correlation map of all genes and miRNAs included in the analysis. The user can visualize the interaractions between multiple miRNAs and multiple targets by considering horizontal or vertical bands in the heatmap.

## Materials and methods

### MoPC Method

The MoPC workflow is reported in Figure 1.

**Fig 1.**
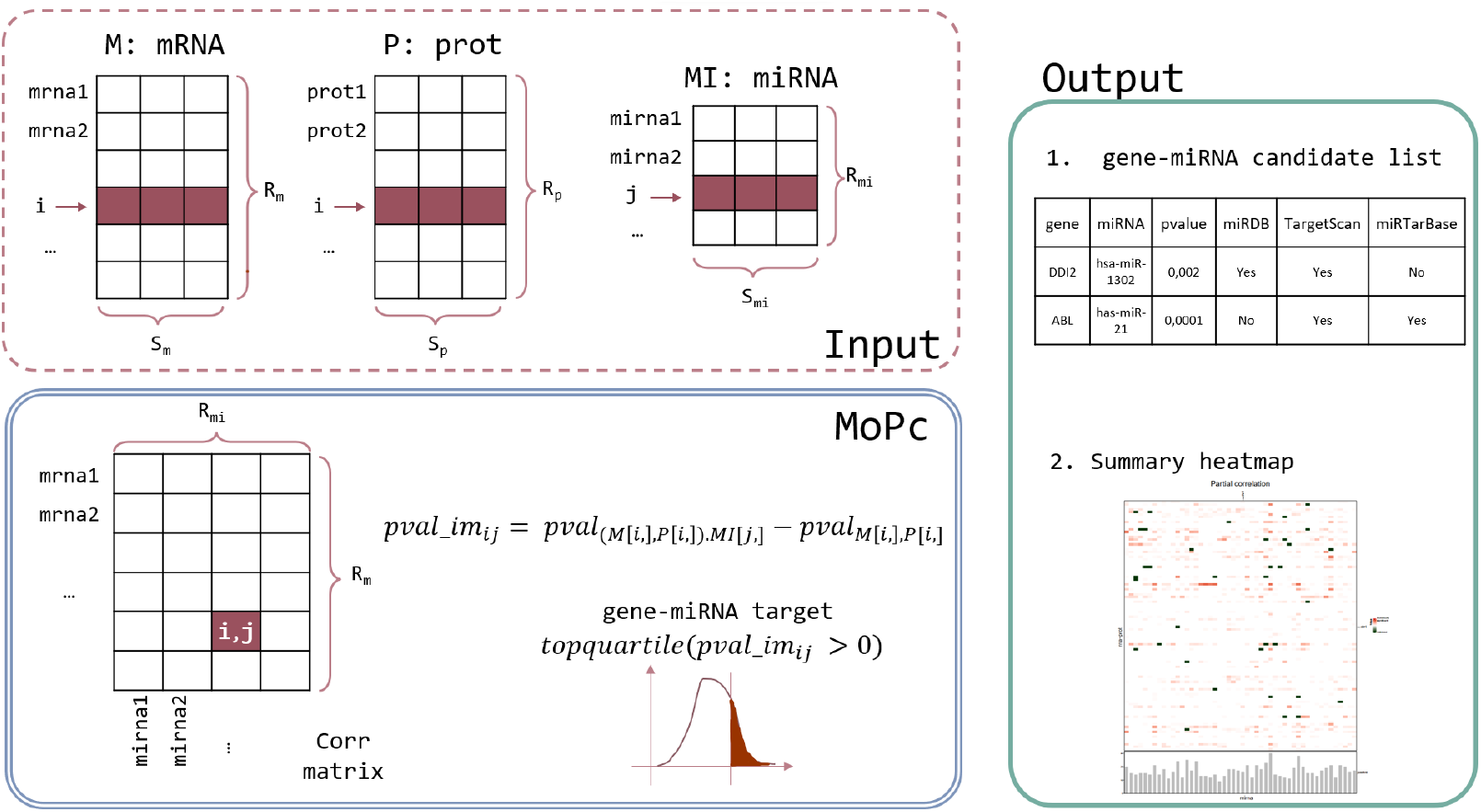
MoPc method in a glance. Input data consist of M, P, and MI matrices representing mRNA, proteomics, and miRNA expression matrices. First, the partial correlation matrix is computed. Note that this matrix has *R*_*m*_ rows corresponding to the number of rows in M and P (genes) and *R*_*mi*_ columns equal to the number of MI rows (miRNAs). Next, the improvement matrix is computed as the difference between the partial correlation p-value and the bivariate correlation p-value. Finally, relevant gene-miRNA interactions are chosen as the top quartile of positive improvement values.

To predict miRNA-target interactions by integrating information at both mRNA and protein level, MoPC calculates the partial correlation of the mRNA and protein levels of a gene conditioned by miRNA expression. Input requires three matrices of mRNA, protein, and miRNA expression denoted respectively as M, P, and MI with the same number of columns n, corresponding to the samples. The M and P matrices must have the same rows IDs corresponding to the genes for which the mRNA and protein levels were measured. Partial correlation *r*_*yx*.*z*_ between mRNA x and protein expression y while accounting for the effect of a given miRNA z is computed as in Eq 1 (*r*_*yx*_ stands for the bivariate correlation between variables *y* and *x*):

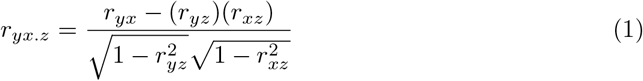

The bivariate Pearson correlation *r*_*yx*_ between mRNA expression x and protein expression y is also computed. Relevant miRNA-target interactions are selected as those for which a lower p-value is observed with partial correlation than that obtained with bivariate correlation. To ensure a large increase in the significance of the p-value, only interactions with an improvement in the p-value in the upper quartile of the distribution are selected.

The MoPc returns two outputs:

1. **miRNA-target significant interactions**. This list reports all miRNA-target candidate interactions based on the partial correlation measure. Additional columns contain the validation information (e.g., if the miRNA-target interaction was previously known in the literature according to three databases: miRDB, TargetScan, and miRTarBase).
2. **summary heatmap**. The heatmap reports genes on the rows and miRNAs on the column ordered according to their chromosomal location. Significant miRNA-targets are reported in red (the more intense the color, the higher the partial correlation estimate). MiRNA-targets both significant and validated in at least one of the three databases (miRDB, TargetScan, and miRTarBase) are reported in dark green. The user can identify miRNAs targeting multiple genes by

visually inspecting horizontal or vertical bands in the heatmap.

The method is freely available as an R package at the following link: https://github.com/martalovino/MoPc

### Data pre-processing

The MoPc method was tested on three TCGA/CPTAC datasets from breast, glioblastoma, and lung cancer studies [19–21].

The breast cancer dataset was obtained from the work of Meryins et al., published in 2016 [19]. The published matrices contain the quantifications of 17814 genes, 7665 proteins, and 332 miRNAs for 77 samples, respectively. The glioblastoma dataset was obtained from the publication of Wang et al. of 2021 [20], including mRNA, protein, and miRNA expression values for 106 tumor samples. The input matrices contain, respectively, 45914 genes, 10998 proteins, and 2883 miRNAs. The lung squamous cell tumor dataset comes from the 2021 publication of Satpathy et al., in which mRNA, proteins, and miRNA expression are quantified [21]. More than 200 samples have been analyzed, of which 201 have all three omics quantifications of our interest. Overall, 21792 genes, 11111 proteins, and 2585 miRNAs are reported.

The three datasets were processed according to the following procedure. These datasets contain missing values, particularly in protein and miRNA expression matrices. For our purposes, proteins and miRNAs with less than 10% of missing values across samples were kept. The remaining missing values were imputed by imposing the median value. Finally, the mRNA e miRNA matrices were log2 transformed.

In all datasets, only the top 50% most highly variable proteins were included, since MoPc requires a certain variance in the expression profiles.

Table 1 shows the datasets size supplied to MoPc.

**Table 1.**
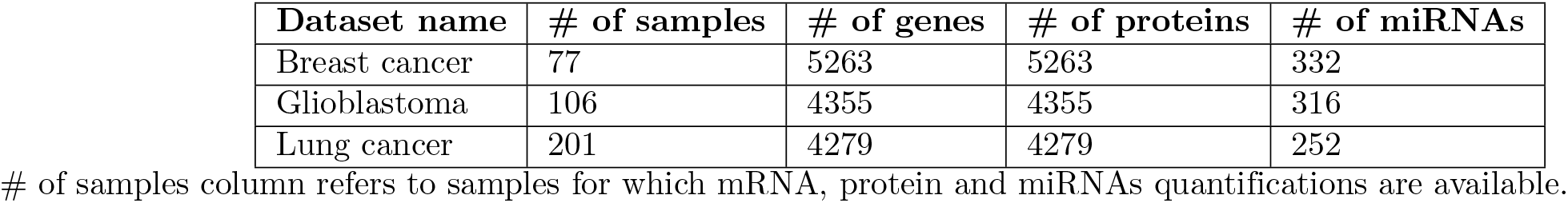
Data dimensions as provided in input to MoPc. For each dataset, the number of samples, quantified mRNA, proteins and miRNAs are reported after the adeguate preprocessing.# of samples column refers to samples for which mRNA, protein and miRNAs quantifications are available.

| Dataset name | # of samples | # of genes | # of proteins | # of miRNAs |
| --- | --- | --- | --- | --- |
| Breast cancer | 77 | 5263 | 5263 | 332 |
| Glioblastoma | 106 | 4355 | 4355 | 316 |
| Lung cancer | 201 | 4279 | 4279 | 252 |

## Results

### MoPC detects biologically relevant regulatory miRNAs in cancer datasets

MoPc was applied to three omic datasets from TCGA/CPTAC projects, namely breast, glioblastoma, and lung cancer. MoPc output returns 1) a table of miRNAs and predicted target genes and 2) a heatmap to visualize the chromosomal position of miRNAs and target genes. In particular, the table listing miRNAs and target genes contains both the estimate value and p-value returned from the partial correlation (the higher the estimate and the lower the FDR adjusted p-value, the more significant the interaction between that gene and that miRNA). In addition, it returns if the gene-miRNA target interaction is already annotated in the three external databases (miRDB, TargetScan, and miRTarBase).

### Computational validation

In order to test the reliability of MoPC predictions, the enrichment in miRDB, TargetScan, and miRTarBase predicted targets was assessed for each of the three cancer datasets, using hypergeometric test. In addition, the same enrichment test was performed for miRNA-target predictions based on the bivariate correlation between mRNA and miRNA expression and between protein and miRNA expression. Pearson correlation with FDR corrected p-values was computed and the miRNA-target pairs with a p-value lower than alpha 0.05 were selected. The same conditions of the partial correlation were used to compute the bivariate correlations to perform a fair comparison. Tables 2, 3, and 4 report for each of the three cancer datasets the percentage of miRNA-targets selected as significant, and the p-value of the hypergeometric test in each of the three databases (miRDB, TargetScan, and miRTarBase) both for the MoPc method and the mRNA-miRNA and protein-miRNA bivariate correlation.

**Table 2.** Validation of results on breast cancer dataset. The first two columns contain the results related to the bivariate correlation between mRNA-miRNA and protein-miRNA. The last column contains the MoPc results.

|  | Bivariate correlation<br>mRNA-miRNA | Bivariate correlation<br>protein-miRNA | MoPc based on<br>partial correlation |
| --- | --- | --- | --- |
| % of predicted miRNA-targets | 2,59% | 0,68% | 17,20% |
| miRDB enrichment | 2.66e-05 | 0.607 | 3.73e-05 |
| TargetScan enrichment | 0.038 | 0.094 | 0.0003 |
| miRTarBase enrichment | 0.0094 | 0.023 | 2.24e-11 |

**Table 3.** Validation of results on glioblastoma dataset. The first two columns contain the results related to the bivariate correlation between mRNA-miRNA and protein-miRNA. The last column contains the MoPc results.

|  | Bivariate correlation<br>mRNA-miRNA | Bivariate correlation<br>protein-miRNA | MoPc based on<br>partial correlation |
| --- | --- | --- | --- |
| % of predicted miRNA-targets | 36,69% | 41,70% | 18,26% |
| miRDB enrichment | <e-70 | <e-70 | 0.999 |
| TargetScan enrichment | <e-70 | <e-70 | 9.48e-23 |
| miRTarBase enrichment | <e-70 | <e-70 | 1.57e-63 |

**Table 4.** Validation of results on lung cancer dataset. The first two columns contain the results related to the bivariate correlation between mRNA-miRNA and protein-miRNA. The last column contains the MoPc results

|  | Bivariate correlation<br>mRNA-miRNA | Bivariate correlation<br>protein-miRNA | MoPc based on<br>partial correlation |
| --- | --- | --- | --- |
| % of predicted miRNA-targets | 61,13% | 59,60% | **18,49%** |
| miRDB enrichment | <**e-70** | <**e-70** | 2.891e-11 |
| TargetScan enrichment | <**e-70** | <**e-70** | 2.973e-06 |
| miRTarBase enrichment | <**e-70** | <**e-70** | 6.68e-27 |

**Table 5.** Breast cancer Validation information.

| miRNA | gene/pathways | known effect | Reference |
| --- | --- | --- | --- |
| hsa-miR-3200-3p |  |  |  |
| hsa-miR-197-3p |  |  |  |
| hsa-miR-505-5p |  |  |  |
| hsa-miR-505-3p |  |  |  |
| hsa-miR-18a-5p |  |  |  |
| hsa-miR-17-5p |  |  |  |
| hsa-miR-577 |  |  |  |
| hsa-miR-135b-5p | LATS2, CDK2, p-YAP | Promotion of cell proliferation and S-G2/M cell cycle progression | [22] |
| hsa-miR-93-3p |  |  |  |
| hsa-miR-93-5p |  |  |  |
| hsa-miR-20a-3p |  |  |  |
| hsa-miR-301a-3p |  |  |  |
| hsa-miR-127-3p |  |  |  |
| hsa-miR-29b-1-5p | Akt3, VEGF, c-MYC | Anti-angiogenesis and anti-tumorigenesis | [22] |
| hsa-miR-15b-3p | Cyclin E1, E2F7 | Anti-proliferative and G1-S cell cycle arrest | [22] |
| hsa-miR-17-3p |  |  |  |

**Table 6.** Glioblastoma Validation information.

| miRNA | gene/pathways | known effect | Reference |
| --- | --- | --- | --- |
| hsa-miR-128-3p | Tumor suppressor | Proliferation, apoptosis, angiogenesis, stemness, radioresistance | [25] |
| hsa-miR-139-5p | In vitro, human | ELTD1, cyclin A, cyclin D1 | [25] |
| hsa-miR-21-5p | oncomiR | Survival, proliferation, apoptosis, migration, invasion, chemoresistance | [25] |
| hsa-miR-21-3p | oncomiR | Survival, proliferation, apoptosis, migration, invasion, chemoresistance | [25] |
| hsa-miR-7-5p | Tumor suppressor | Survival, proliferation, apoptosis, invasion, angiogenesis | [25] |
| hsa-miR-1249-3p |  |  |  |
| hsa-miR-155-5p | In vitro, human | MAPK13, MAPK14 | [25] |
| hsa-miR-124-3p | Cdk6, PPPR13L, SNAI2/GJIC | miR-124-3p inhibits glioblastoma cell growth in part by diminishing the protein expression of cdk6, leading to cell cycle arrest at the G0/G1 phase. | [25] |
| hsa-miR-874-3p |  |  |  |
| hsa-miR-15b-5p | In vitro, human | miR-15b inhibited the proliferation, cell cycle arrest, and invasion. | [25] |
| hsa-miR-769-5p |  |  |  |
| hsa-miR-132-5p | In vitro, human | miR-132 promoted the proliferation and sphere formation of cells. | [25] |

**Table 7.** Lung cancer Validation information.

| miRNA | gene/pathways | known effect | Reference |
| --- | --- | --- | --- |
| hsa-miR-182-5p |  |  |  |
| hsa-miR-26b-5p |  |  |  |
| hsa-let-7a-5p | LIN28 | Chemo sensitive | [24] |
| hsa-miR-26a-5p |  |  |  |
| hsa-miR-183-5p |  |  |  |
| hsa-let-7f-5p | LIN28 | Chemo sensitive | [24] |
| hsa-miR-30a-3p | PI3K-SIAH2 | EGFR TKI-resistant | [24] |
| hsa-miR-125a-5p |  |  |  |
| hsa-miR-205-5p | PTEN, Mcl-1 and Survivin | Chemo resistant | [24] |
| hsa-miR-126-3p | Akt and ERK pathways | Chemo sensitiveEGFR TKI-sensitive | [24] |
| hsa-miR-7-5p | EGFR | Chemo sensitive | [24] |
| hsa-miR-9-3p |  |  |  |

### MiRNA-targets visualization map

MoPc returns a heatmap where miRNAs and predicted targets are arranged according to their chromosomal position. The heatmap represents in red the miRNA-targets significantly predicted according to the MoPc partial correlation analysis and in dark green, the miRNA-targets statistically predicted and validated in at least one of the three databases (miRDB, TargetScan, and miRTarBase). The user can visually explore the presence of miRNAs that regulate multiple genes and vice-versa (the same gene regulated by multiple miRNAs). Figures S1, S2, S3 show breast, glioblastoma, and lung cancer heatmaps, respectively.

## Discussion

The goal of MoPc is to predict relevant miRNA-targets integrating information from miRNA, mRNA and proteome profiles. The predicted targets are validated with three reference databases (miRDB, TargetScan, and miRTarBase), both for the MoPc method and for the bivariate correlation between mRNA-miRNA and protein-miRNA. As reported in Tables 2, 3, and 4, MoPc predictions are enriched in independent database targets for the three cancer datasets analyzed. Except for the enrichment of glioblastoma in miRDB, all enrichments are statistically significant. The miRNA-target pairs predicted from the bivariate mRNA-miRNA and protein-miRNA correlation are also statistically enriched for each cancer dataset and database, unless for breast cancer in the bivariate protein-miRNA correlation in the mirDB database.

An essential aspect to compare is the total number of miRNA-target interactions predicted by MoPc and bivariate mRNA-miRNA and protein-miRNA correlation. Analyses of glioblastoma and lung cancer datasets benefit from the results obtained through MoPc. Indeed, MoPc returns 17.20%, 18.26%, and 18.49% of miRNA-target interactions for breast, glioblastoma, and lung cancer. Although the bivariate mRNA-miRNA and protein-miRNA correlation are generally more enriched in the three external target databases, they return a higher percentage of miRNA-target correlated pairs, typically greater than 36% for both mRNA-miRNA and protein-miRNA. This aspect is of uttermost importance, as the number of investigable gene-miRNA pairs is not clinically useful if it is close to 30-60% of all gene-miRNA pairs in the input dataset. MoPc proves to be an effective method in identifying key miRNAs within a dataset, reducing the number of targets while maintaining the statistical significance of the enrichment. After the MoPc computational validation in the three external databases, we focused on specific miRNAs known in the literature to be associated with specific diseases. Indeed, the enrichment in miRDB, TargetScan, and miRTarBase verifies that the miRNA-target interaction is known or predicted in the literature. However, such enrichment does not consider whether the predicted miRNAs are key in the specific disease (breast, glioblastoma, lung cancer). Therefore we compared MoPc miRNA-target pairs with disease-associated miRNA in the literature [22–24].

Regarding breast cancer, MoPc detects the main miRNAs involved in the cancer genesis, including miR-126 and miR-140 (with anti-angiogenesis and anti-tumorigenesis function) and miR-200b and miR-200c (promoting metastasis and invasion). Furthermore, among the miRNAs known in glioblastoma, MoPc successfully identifies the tumor suppressors miR-7 and miR-128 and the oncomiRs miR-21 and miR-93, responsible for survival, proliferation, apoptosis, and drug resistance. For lung cancer, the miRNAs known and identified by MoPc are miR-7 and miR-21 (chemosensitive and chemoresistant, respectively) and miR-200c and miR-223 with EGFR-TKI and ALK-TKI sensitivity. The Tables below show the top 5 percentile of miRNAs with the highest significant number of targets found by MoPc and their associated disease.

## Conclusion

Identifying key miRNAs is of fundamental importance for the early diagnosis and treatment of many diseases, including cancer. However, in the literature, various tools deal with providing a list of relevant miRNAs without exploiting the information coming from the expression of proteins. This paper presents MoPc, a novel method for predicting key miRNA-target pairs by exploiting the computation of the partial correlation between mRNA and protein expression conditioned by miRNA expression. The method is applied to three TCGA/CPTAC datasets: breast, glioblastoma, and lung cancer. The results show enrichment for each dataset in the three databases, miRDB, TargetScan, and miRTarBase, except for breast cancer in miRDB. Furthermore, MoPc successfully identifies key miRNAs in each cancer type. Finally, the user can inspect the results both through the list of significant miRNA-gene pairs and graphically through the generated heatmap, in which genes and miRNAs are chromosomally ordered. In the future, this approach can be extended to other types of omics data, given that the underlying biological phenomenon can be detected through partial correlation. the partial correlation p-value and the bivariate correlation p-value. Finally, relevant gene-miRNA interactions are chosen as the top quartile of positive improvement values.

## Supporting information

**S1 Fig. Breast cancer summary heatmap**. The heatmap reports in red the miRNA-genes pairs resulting significantly with the MoPc analysis. In dark green, the miRNA-genes pairs are statistically significant in the input dataset and validated in at least one of the three databases (miRDB, TargetScan, and miRTarBase). Genes are reported on the rows, and miRNAs on the columns. Both genes and miRNAs are chromosomally ordered.

**S2 Fig. Glioblastoma summary heatmap**. The heatmap reports in red the miRNA-genes pairs resulting significantly with the MoPc analysis. In dark green, the miRNA-genes pairs are statistically significant in the input dataset and validated in at least one of the three databases (miRDB, TargetScan, and miRTarBase). Genes are reported on the rows, and miRNAs on the columns. Both genes and miRNAs are chromosomally ordered.

**S3 Fig. Lung cancer summary heatmap**. The heatmap reports in red the miRNA-genes pairs resulting significantly with the MoPc analysis. In dark green, the miRNA-genes pairs are statistically significant in the input dataset and validated in at least one of the three databases (miRDB, TargetScan, and miRTarBase). Genes are reported on the rows, and miRNAs on the columns. Both genes and miRNAs are chromosomally ordered.

